# Contributions by N-terminal Domains to NMDA Receptor Currents

**DOI:** 10.1101/2020.08.21.261388

**Authors:** Stacy A. Amico-Ruvio, Meaghan A. Paganelli, Jamie A. Abbott, Jason M. Myers, Eileen M Kasperek, Gary I. Iacobucci, Gabriela K. Popescu

**Affiliations:** From the Department of Biochemistry and the State University of New York, Buffalo, New York 14214; Neuroscience Program University at Buffalo, State University of New York, Buffalo, New York 14214

**Keywords:** Ionotropic glutamate receptors, NMDA receptors, channel activation, single-molecule bio-physics, allosteric regulation, receptor structure-function

## Abstract

To investigate the role of the N-terminal domains (NTDs) in NMDA receptor signaling we used kinetic analyses of one-channel currents and compared the reaction mechanism of recombinant wild-type GluN1/GluN2A and GluN1/GluN2B receptors with those observed for NDT-lacking receptors. We found that truncated receptors maintained the fundamental gating mechanism characteristic of NMDA receptors, which includes a multi-state activation sequence, desensitization steps, and mode transitions. This result establishes that none of the functionally-defined NMDA receptor activation events require the NTD. Notably, receptors that lacked the entire NTD layer retained isoform-specific kinetics. Together with previous reports, these results demonstrate that the entire gating machinery of NMDA receptors resides within a core domain that contains the ligand-binding and the channel-forming transmembrane domains, whereas the NTD and C-terminal layers serve modulatory functions, exclusively.

## INTRODUCYION

NMDA receptors are excitatory neurotransmitter receptors with prominent and vital functions in the mammalian central nervous system (CNS). The electrical signals generated by NMDA receptors result from complex reaction mechanisms that include agonist-binding reactions and dynamic isomerizations that accomplish gate opening, channel desensitization, and mode transitions (1–3). Despite increasing knowledge of NMDA receptor architectures, the specific structures that support activation, desensitization and moding remain under investigation (4–9). Mammalian NMDA receptors are the largest and most complex of neurotransmitter-gated channels (10). Like all 18 genes in the ionotropic glutamate receptor (iGluR) family, the GluN genes that make the NMDA receptor family have mosaic structures and encode four distinct semi-autonomous domains that speak of evolutionary kinship to simpler proteins (11–15). The two extracellular domains, the N-terminal (NTD) and the membrane proximal ligandbinding (LBD) domains, share homology with the bacterial periplasmic binding-proteins (PBP) LAOBP and QBP, respectively; the transmembrane domain (TMD) resembles, in part, the pores of primitive cation-selective channels; and lastly, the C-terminal domains (CTDs), which reside intracellularly and represent the most variable region of iGluR subunits, are unrelated to known proteins(10).

Several functional behaviors pertinent to intact receptors have been traced to distinct structural domains, which maintain overall structural integrity and function even when experimentally separated from the full protein. The fundamental aspects of channel activation, *i. e*. agonist binding sites and ligand-controlled gate, have been mapped within LBDs (13, 16–18) and TMDs (15,19,20), respectively, with increasingly prominent roles in coupling assigned to stretches of residues that link the LBD with the TMD (21–23). Far less is known about the structures that support desensitization and moding, leaving constitutive steps of the activation process structurally unspecified.

Consistent with agonist-binding and ionic flux being mediated by LBD and TMD, respectively, neither the NTDs nor the CTDs are essential to agonist-dependent activation (24–30). In previous work, we showed that although CTDs of NMDA receptors modulate channel conductance and open probabilities, CTD-lacking receptors display all functional properties of wild-type receptors including desensitization and modal gating, consistent with a purely modulatory role in channel function (31). The work reported here examines the roles of NTDs in the NMDA receptor reaction mechanism.

Recombinant receptors lacking NTDs insert in the plasma membrane and retain glutamate-gated ionotropy, even when they have reduced surface expression, modified pharmacology, and altered kinetics (24–27,29). In addition, glutamate-activated channels that naturally lack NTDs, are observed in birds (KA-BP) and plants (GluR0), suggesting that the NTDs of mammalian receptors have been appended to an ancestral ionotropic receptor subsequent to the separation of these evolutionary branches (32,33). Within the iGluR family, NTDs host molecular determinants of tetrameric assembly, membrane targeting, and protein-protein interactions. Uniquely to NMDA receptors, NTDs contain several binding sites for endogenous and synthetic effectors, thus assigning functions onto NTD attachments (10).

Glutamatergic NMDA receptors assemble as obligate hetero-tetramers of GluN1 (N1) and GluN2 (N2) subunits. N1 protomers occur as multiple splice variants of a single gene product, with alternative splicing of exon 5 producing two types of NTD modules (a and b); N2 subunits are expressed from four separate genes (A - D), whose NTDs share ~ 30% homology. These structural variations of the NTD modules confer substantial functional and pharmacologic diversity onto the family, and represent therapeutic targets for several neuropsychiatric disorders (34).

Several NMDA receptor allosteric modulators act through residues located on the NTDs of N1 or/and N2 subunits, including protons, polyamines, zinc, and phenyl-ethanol-amines (e. *g*. ifenprodil). The current hypothesis for the action of these pharmacologic agents is that by binding at the interface formed by mobile domains they change the time course of the activation reaction and therefore of the receptor-generated currents. Similarly, isoform-specific structures of the NTDs have been proposed to be a major locus for subunit-specific kinetic differences between NMDA receptor isoforms (35,36).

Yuan et al. (29) studied receptors lacking the NTD of specific N2 subunits (N1/N2A^Δ^, N1/N2B^Δ^, N1/N2C^Δ^ and N2/N2D^Δ^) as well as chimeric receptors whose 2A and 2D subunits had swapped NTDs. Their results uncovered that the NTDs of N2 subunits have no influence on channel conductance but by controlling the time spent by receptors in closed conformations (MCT) they also control important kinetic aspects of the channel response such as open probability (Po) and deactivation time course (τ_d_) (29).

Here, we extend these studies to the entire NTD domain and investigate specifically the mechanisms by which the absence of NTDs affects activation, desensitization, and moding in NMDA receptors. We show that NTD-lacking NMDA receptors: 1) retain characteristic functional properties including agonist-dependent gating, multi-step activation sequence, two microscopic desensitization steps (responsible for bursting behavior), and modal gating; 2) the modulatory role of NTDs is contributed asymmetrically by N1 and N2 subunits; and 3) 2A- and 2B-containing NMDA receptors maintain subunit specific kinetics even in the absence of NTDs.

## RESULTS

### Macroscopic kinetics and zinc sensitivity of NTD-lacking N1/N2A receptors

To delineate contributions by NTDs to the NMDA receptor activation mechanism, we set up to examine current responses from recombinant NMDA receptors that lacked NTD modules of N1 (N1^Δ^), 2A (2A^Δ^), and/or 2B (2B^Δ^) subunits (**Figure 1A, B**). We co-expressed wild-type or NTD-truncated subunits in HEK293 cells in the following combinations: N1^Δ^/N2, N1/N2^Δ^, or N1^Δ^/N2^Δ^, for either 2A or 2B isoforms. Before launching into recording microscopic responses, we aimed to validate functional expression of mutated receptors.

**Figure 1:**
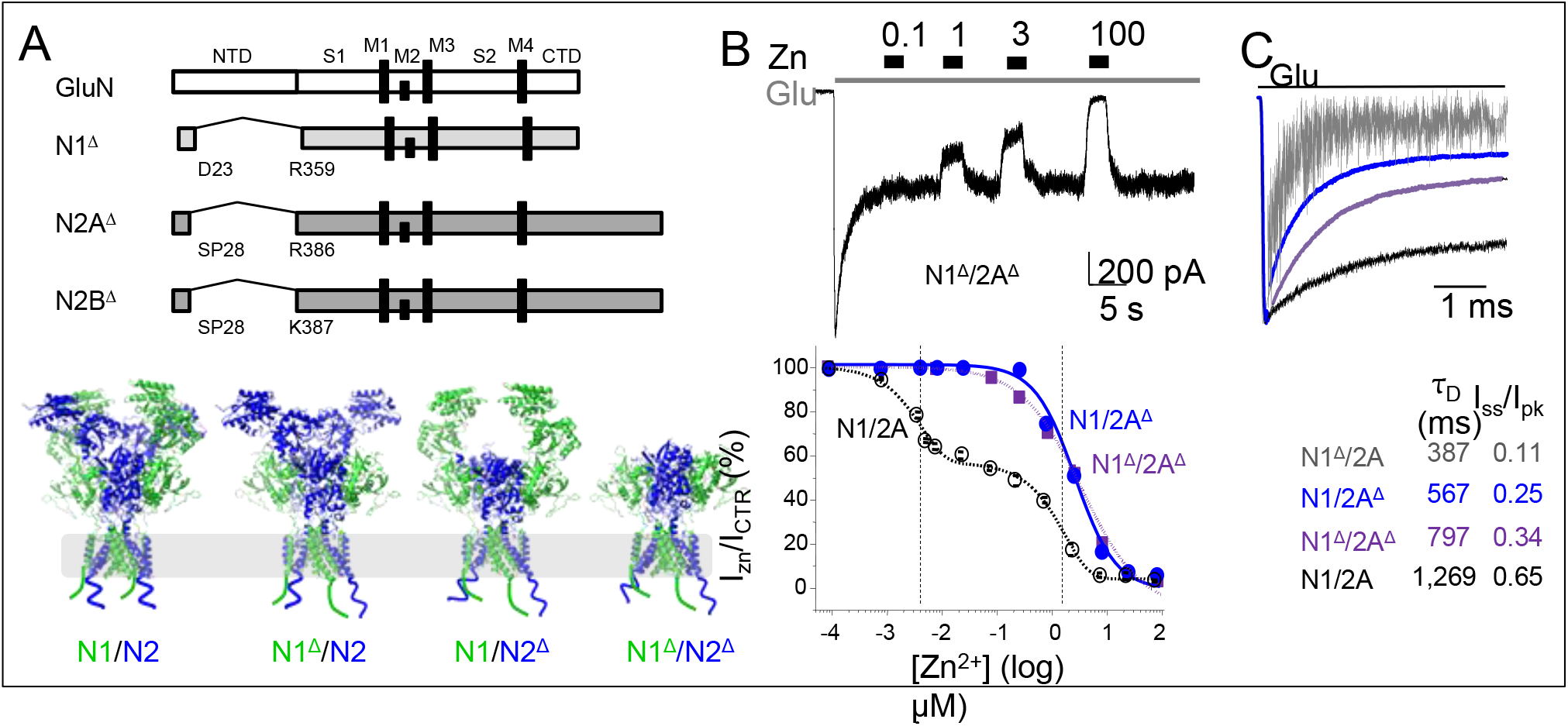
Properties of NTD-lackmg NMDA receptors. *A*, Diagram of the GluN subunit constructs used in this study (top) and hypothetical architectures of the receptors examined based on PDB 4PE5, and schematic of CTD tails of unknown structure (not at scale) (bottom). ***B***, Top, Whole-cell current trace elicited from cells expressing N1^Δ^/2A^Δ^ receptors with a sustained pulse of glutamate (1 mM, and glycine continually present at 0.1 mM), and zinc applications as indicated (in μM). *Bottom*, Zinc-concentration dependent decrease in steady-state current levels (I_zn_/I_CTR_) for wild-type N1/N2A, N1^Δ^/2A^Δ^, and N1/2A^Δ^ receptors. Zinc concentrations that produced half-maximal inhibition (IC_50_) are marked with vertical broken lines for both the high-affinity (nM) and low-affinity (μM) sites. ***C***, Whole-cell current recordings (top) and desensitization kinetics (bottom) as time course (τ_D_) and extent (I_ss_/I_pk_) for wild-type N1/N2A and three truncated receptors.

As a measure of N2A NTD function, we examined glutamate-elicited responses and tested their sensitivity to zinc inhibition (37,38). We recorded wholecell currents during applications of glutamate (5-s, 1 mM) in the continuous presence of glycine (0.1 mM), and applied increasing concentrations of zinc to the steady state portion of the current. As expected for receptors lacking the NTD of N2A, N1/2A^Δ^ and N1^Δ^/2A^Δ^ receptors lost their high-affinity sensitivity to Zn^2+^ but retained low-affinity Zn^2+^ sensitivity (**Figure 1B**) (39–41).

Our results also showed drastic changes in macroscopic response kinetics. Consistent with previous reports, N1/N2A receptors lacking NTDs produced macroscopic currents that desensitized faster, as measured by shorter decay time constants (τ_D_) and deeper, as measured by a reduction in the relative amplitudes of the steady-state and peak currents (I_ss_/I_pk_) (**Figure 1C**). The most dramatic effect occurred for receptors lacking the NTDs of N1 subunits alone, whereas receptors lacking only the NTDs of N2A subunits and those lacking the entire NTD layer showed intermediate phenotypes.

In keeping with a proposed role for the N1-NTD in receptor assembly and efficient surface expression (25), whole-cell currents from cells expressing N1^Δ^/2A receptors were ~20-fold smaller than those obtained from cells expressing wild-type receptors (96 ± 44 pA, n = 9 vs. 2,340 ± 382 pA, n = 10). In contrast, cells expressing receptors lacking the NTD of 2A subunits, N1/2A^Δ^ and N1^Δ^/2A^Δ^, produced currents of comparable amplitudes: 1,010 ± 435 pA (n = 6) and 1,850 ± 294 pA (n = 7), respectively (p > 0.05, Student’s t-test).

Smaller whole-cell currents may reflect reduced surface expression but also decreased single-channel conductance and/or alterations in activation kinetics. We investigated these possibilities in greater detail, with single-channel current recordings.

### Unitary currents from NTD-lacking receptors display characteristic NMDA receptor patterns

We recorded single-channel currents from cell-attached patches containing only one active receptor and measured their unitary current amplitudes (**Figure 2).** Regardless of which NTD domain was truncated, unitary amplitudes were similar with those measured for wild-type receptors for both N1/N2A and N1/N2B receptors (**Table 1**). This observation is consistent with previous reports that the NTD of N2 subunits has no effect on unitary conductance (29). In addition, they demonstrate that channel conductance is independent of the presence of the N1 NTD, and of the presence of the entire NTD layer.

**Figure 2:**
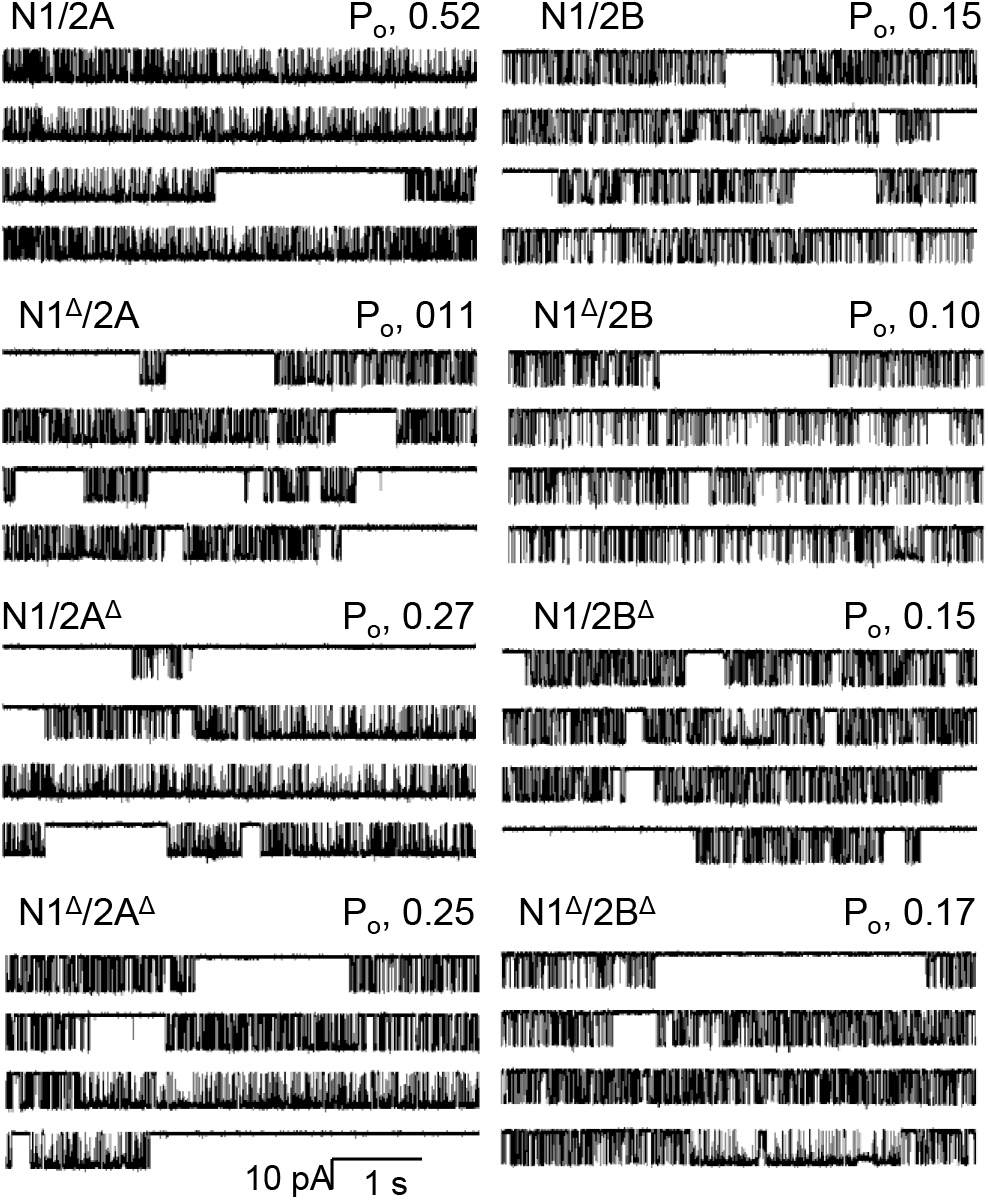
Effects of NTD truncations on NMDA receptor single-channel currents. Continuous traces were recorded from cell-attached patches that contained one active channel exposed to 1 mM Glu, and 0.1 mM Gly. Each panel illustrates 20-s from a minutes-long recording (5-s/trace, 1 kHz filter) for the indicated receptor.

**Table 1:** Kinetic parameters of single NMDA receptors.

| receptor | n | amp<br>(pA) | P<sub>o</sub> | MCT<br>(ms) | MOT<br>(ms) | Duration<br>(min) | Events<br>(X10<sup>6</sup>) |
| --- | --- | --- | --- | --- | --- | --- | --- |
| N1/2A | 13 | 8.7 ± 0.3 | 0.52 ± 0.03 | 6 ± 1 | 6.7 ± 0.4 | 435 | 3.8 |
| N1<sup>Δ</sup>/2A | 7 | 9.9 ± 0.7 | 0.10 ± 0.03* | 60 ± 9* | 5.6 ± 0.7 | 367 | 0.8 |
| N1/2A<sup>Δ</sup> | 10 | 10.4 ± 0.6 | 0.27 ± 0.05* | 27 ± 4* | 8.5 ± 1.2 | 314 | 1.1 |
| N1<sup>Δ</sup>/2A<sup>Δ</sup> | 11 | 9.7 ± 0.7 | 0.25 ± 0.04* | 27 ± 4* | 7.1 ± 0.6 | 317 | 1.2 |
| N1/2B | 16 | 10.6 ± 0.4 | 0.15 ± 0.03 | 36 ± 5 | 4.6 ± 0.5 | 552 | 1.6 |
| N1<sup>Δ</sup>/2B | 8 | 10.8 ± 0.2 | 0.10 ± 0.02 | 39 ± 11 | 2.8 ± 0.2* | 271 | 0.9 |
| N1/2B<sup>Δ</sup> | 5 | 9.8 ± 0.4 | 0.15 ± 0.04 | 31 ± 5 | 4.8 ± 0.6 | 387 | 1.8 |
| N1<sup>Δ</sup>/2B<sup>Δ</sup> | 6 | 11.7 ± 0.6 | 0.17 ± 0.03 | 23 ± 5 | 3.7 ± 0.4 | 303 | 1.8 |

\* indicates significant differences relative to wild-type ( $p < 0.05$ , Student's $t$ -test); higher, *red* or lower, *blue*.

Next, we examined the gating kinetics of NTD-lacking receptors, as observed in one-channel records. All receptors used in this study gated with complex single-channel behaviors that included bursts of channel openings and closings (42,43), separated by long closures designated as desensitized intervals (**Figure 2**). On first pass, this observation indicates that the NTD is not responsible for the characteristic bursting and desensitizing features of single NMDA receptors.

Further, kinetic modeling of these data sets, described in more depth below, showed that single-channel traces were adequately described with models that had five closed components and up to four open components, as previously reported for wildtype 2A and 2B receptors (**Figures 3**, **4**, and **Table 2**) (44–46). This observation indicates that truncated receptors maintained a multi-state opening sequence (47–49), two desensitized states (44,47), and modal gating (1,2,48).

**Figure 3:**
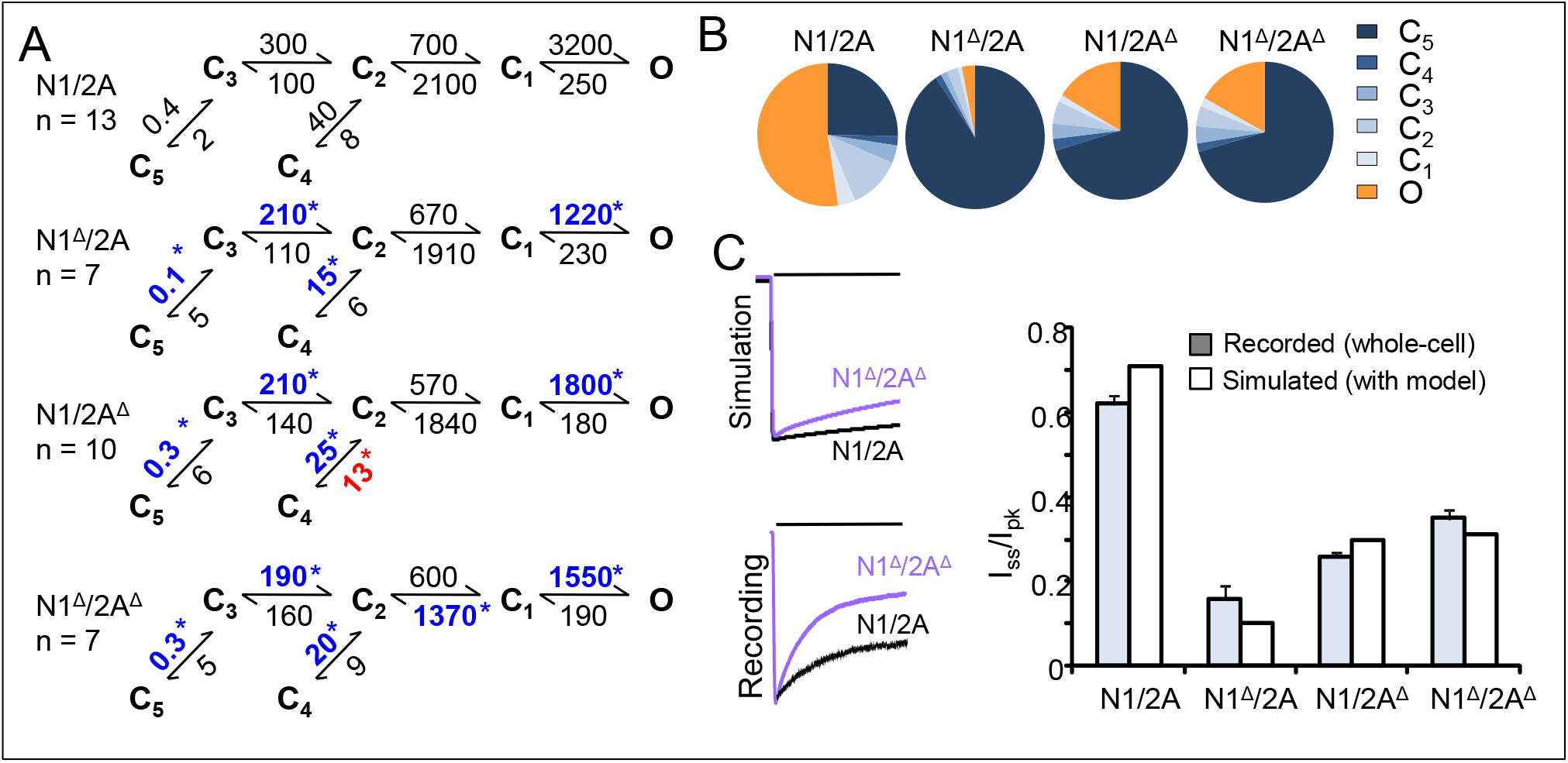
Effects of NTD truncations on N1/N2A receptor gating kinetics. *A*, Reaction mechanisms were estimated with the illustrated models from direct fits to single-channel data. All states represent receptor conformations fully liganded with glutamate and glycine: C, non-conductive; O, conductive. Rate constants are given above the respective arrows as average values for each dataset (s^−1^). * indicates values that are different (faster, *red* or slower, *blue*) relative to N1/2A (*p* < 0.05 Student’s *t*-test). *B*, State occupancies calculated from the reaction mechanisms in *A. C*, Macroscopic responses simulated with the models in *A* recapitulate the extent of desensitization (I_ss_/I_pk_) observed in experimentally recorded traces (as in Figure 1D).

**Figure 4:**
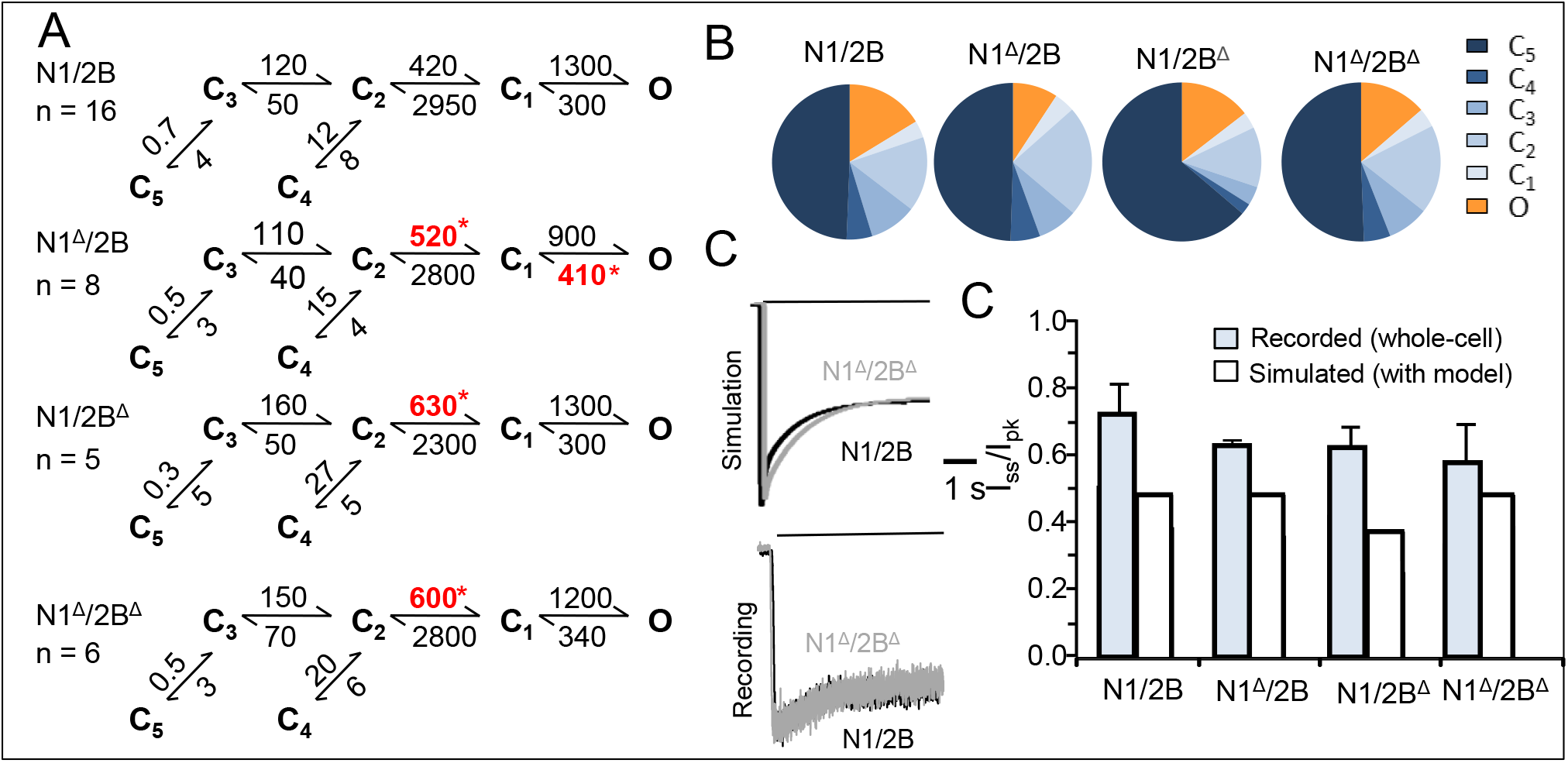
Effect of NTD truncations on the kinetic reaction mechanisms of 2B-type NMDA receptors. Reaction mechanisms estimated from fitting a 5C1O model to single-channel data. All states represent receptor conformations fully liganded with glutamate and glycine; C, non-conductive; O, conductive. Rate constants (s^−1^) are given as average values for each data set. * indicates significant differences (faster, *red* or slower, *blue*) relative to wild-type receptors (*p* <0.05, Student’s *t*-test). *B*, State occupancies calculated from the reaction mechanisms in *A. C*, Macroscopic responses simulated with the models in *A* recapitulate the extent of desensitization (I_ss_/I_pk_) observed in experimentally recorded traces.

**Figure 5:**
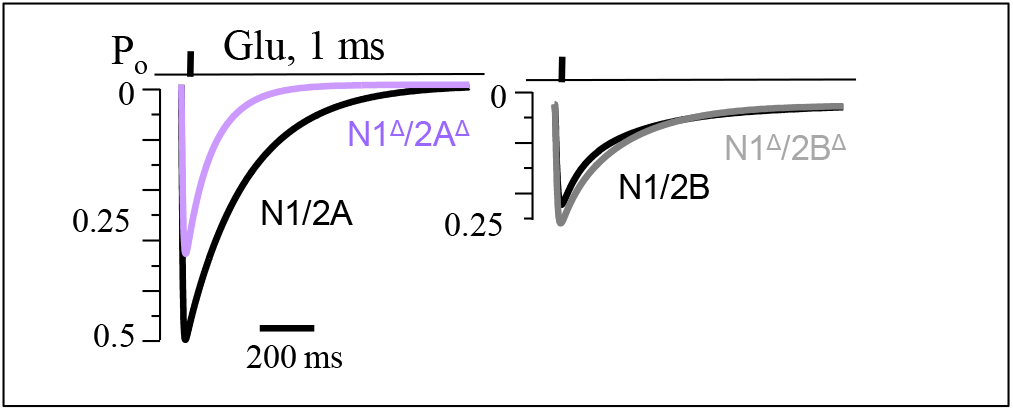
Effects of NTD truncations on synaptic-like responses. Macroscopic responses to one 1-ms glutamate pulse were simulated with the kinetic models for the indicated NMDA receptors.

**Table 2:**
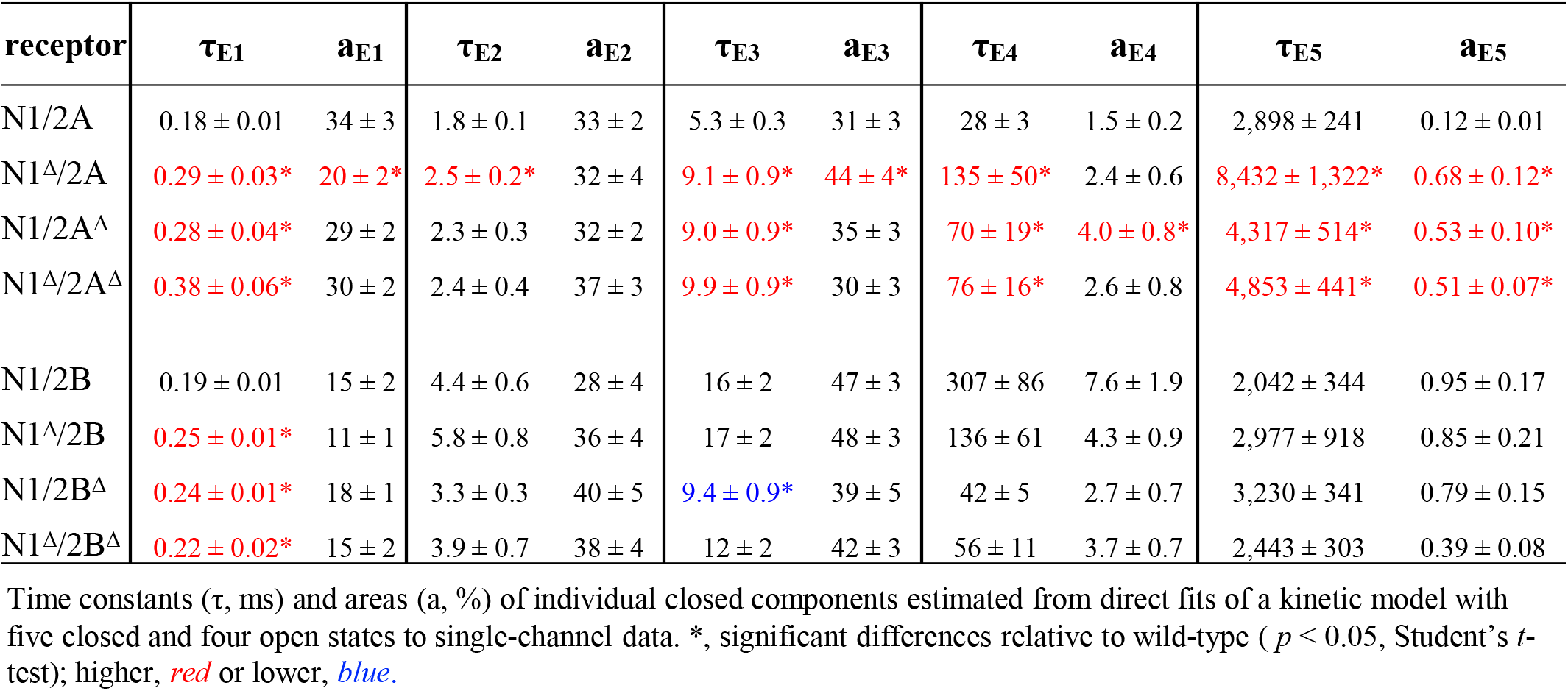
Effects of NTD truncation on individual closed components.

In summary, truncation of the NTDs of either one or both subunits had no effect on the number of closed and open components present in these data, suggesting that the NTD itself does not change the basic kinetic scheme of NMDA receptor activation.

### Subtype-dependent contributions of NTD to Po

In addition to information about unitary amplitudes, bursting behavior, and bursting structure, unitary currents, especially when recorded from a single active molecule as in this study, inform about the absolute open probability of the channel examined.

Truncating NTDs of N1/N2A receptors, whether in only one subunit or both, decreased the channels’ open probability (P_o_) by increasing closed durations, and had no effect on open durations (**Table 1).** In contrast, truncating NTDs of N1/N2B receptors had little if any effect on P_o_ (**Table 1**). Regardless of the extensive truncations, all three of the N1/N2B constructs examined had largely wild-type behaviors.

### In N1/N2A receptors, NTDs increase Po by destabilizing desensitized states

To elucidate the role of NTDs in channel gating we fit our single-channel data for wild-type and NTD-truncated receptors with kinetic multi-state models and estimated rate constants for the transitions included in the model. As noted previously, all records were well described with five closed and two to four open components. Multiple open components are indicative of modal gating, with each recording capturing anywhere from one mode (two open states) and up to three modes (four open states) (1,2,46,48,50). Due to the complexity of the reaction mechanism, for ease of comparison across recordings, and given that open times were unaffected by NTD truncations, we used for subsequent measurements a simplified kinetic scheme where a single open state, aggregated all open states observed in each recording (**Figures 3, 4**). We used this model to identify the microscopic transitions whose kinetics were affected by truncation of the NTDs.

For N1/N2A receptors, NTD deletions affected several rate constants in the reaction mechanism. Consistently, for all three NTD-truncated N1/N2A receptors, rate constants returning receptors from either desensitized state into the active gating sequence (k_53_, k_42_) were ~2-fold slower (**Figure 3A**). Together with slower rates along the activation sequence (k_32_, k_1O_), these changes increased substantially the occupancy of desensitized states (**Figure 3B**).

To validate these kinetic models, we used them to simulate macroscopic currents, and compared these traces with the experimentally recorded currents (**Figure 3C**). Simulated currents had kinetics (I_ss_/I_pk_) that matched well the recorded currents indicating that the reaction mechanisms deduced from kinetic modeling of one-channel recordings reflects well the observed behaviors of assemble responses. This validation strengthens the conclusion that when present, NTDs of N1/N2A receptors increase channel P_o_ by accelerating the opening reaction and destabilizing desensitized states.

### In N1/N2B receptors, truncation of NTDs has negligible effect on gating kinetics and P_o_

Unexpectedly, in N1/N2B receptors, truncation of NTD in either subunit, or of the entire NTD layer had only a marginal effect on the channel’s reaction mechanism (**Figure 4A**, **Tables 1, 2**). None of the examined truncations produced significant effects on channel P_o_ or closed durations (MCT). N1/N2B receptors lacking the N1 NTD, had a small but statistically significant reduction in mean open durations (from 4.6 ± 0.5 ms to 2.8 ± 0.2 ms, p < 0.05, Student’s t-test), which resulted in a slight reduction in channel P_o_ from (0.15 ± 0.03 to 0.10 ± 0.02, p > 0.05, Student’s t-test). However, due to the well-documented variability in closed times for this receptor type (46), the reduction in P_o_ was not statistically significant. Instead, the model predicts a redistribution of receptors across closed states (**Figure 4B**), resulting in increased occupancy of the longest desensitized state (C5), as observed for NTD truncations in N1/N2A receptors (**Figure 3B**).

We were surprised by the minimal effect of NTD truncations on N1/N2B receptors kinetics and we aimed to test the model by comparing its macroscopic predictions with experimentally recorded currents. We used the model to simulate ensemble responses to glutamate (1 mM) and compared these traces with experimentally recorded whole-cell responses from cells expressing each truncated receptor. Results show that as predicted by the model, NTD-lacking N1/N2B receptors had minimal changes in their macroscopic desensitization kinetics (**Figure 4C**).

Together, these observations and analyses indicate that when present, NTDs of N1/N2B receptors have only small effects on overall channel Po, but produce a redistribution across closed states, and this is most prominent for the N2B NTD.

### NTD-truncated receptors maintain isoform specific kinetics

In CNS, the majority of excitatory synapses express N2A- and N2B-containing receptors whose ratios are dynamic and among other factors depend on the developmental stage of the animal, and the maturity and strength of each synapse (51–54). In practice, the exact molecular composition of synaptic NMDA receptors is difficult to ascertain. Instead the decay time of the excitatory synaptic current (τ_d_) is used as a proxy for molecular composition, based on the markedly slower (~3-4-fold) decay time of recombinant N2B- and N2A-containing receptors (55,56). Given the stronger sequence homology in LBD and NTD domains, these characteristic and biologically relevant differences in decay time have been attributed in large part to differences in the NTDs of the two isoforms.

We used the models obtained from single-channel currents (**Figure 2**) and validated with whole-cell recordings (**Figures 3, 4**) to simulate synaptic-like responses following stimulation with a brief (1 ms) pulse of glutamate (1 mM). Results show that although NTD-lacking N1/N2A receptors respond with lower peak Po, which indeed makes them more similar to the less active N1/N2B receptors, they also have faster deactivation kinetics, which makes them even more different than the slow decaying N1/N2B currents. Importantly, in practice it is difficult to measure the peak Po, which has been measured for only a handful of preparations. Instead, experimental recordings are normalized to peak and evaluated in terms of their deactivation time. Our results suggest that NMDA receptors with NTD truncations maintain isoform-specific kinetics.

## DISCUSSION

We used one-channel current recordings and kinetic modeling of NMDA receptors lacking NTD modules of N1, N2B or N2A subunits or the entire NTD layer to quantify for the first time the role of NTD domains in NMDA receptor gating. We report three main conclusions. First, truncated receptors retained all the gating features characteristic of NMDA receptors, *i.e*. multi state activation, two desensitized states and three kinetic modes. Second, NTD truncations reduced the gating kinetics of N1/N2A and had minimal effect on the gating of N1/N2B receptors. Third, receptors lacking the entire NTD layer retained isoform-specific kinetic differences.

Definitive evidence indicates that agonist-dependent gating of NMDA receptors require functional LBDs and TMD and their cooperation (4, 5, 7, 57–59). Further, CTD domains modulate channel conductance and channel open probability, and mediate effects on conductance, calcium-permeability, and gating by intracellular effectors (31,60,61). However, CTD-lacking receptors maintain desensitization, the ability to go through modal transitions, and isoform-specific kinetics. The results we present here, complete the picture by showing that NTD-lacking receptors have the necessary machinery to perform all the basic functions of ligand-dependent gating, desensitization, and moding. The implication is that like ligand-dependent gating, the structural determinants of desensitization and moding also reside within the structural core represented by LBDs, NTD and their coupling.

This knowledge is important because the large size of NTD and CTD layers of the NMDA receptors, which represent more than half their mass, precludes their structural examination in intact form. In fact, the most comprehensive structural models reported to date for tetrameric NMDA receptors describe CTD-lacking receptors (4–8,58,59,62). Further, tetrameric receptors that lack both the NTD and the CTD layers, represent more amenable targets for high resolution structural studies (7). These are also feasible for full-atom computational approaches aimed to map the temporal aspects of their activation mechanism. The results we present here, together with previous reports, indicate that such minimal, or core receptors maintain the essential components of ligand-dependent gating, desensitization, and moding, and therefore they represent valid preparations for in-depth structural and molecular dynamics studies. Importantly, they also retain characteristic isoform-dependent kinetics, whose yet-unknown underpinning control important biological functions.

In summary, our results support the value of minimal, or core, NMDA receptor proteins that lack NTD and LBD layers as subjects for future high-resolution structural investigations and for computational approaches to map the dynamic structural changes that make the NMDA receptor activation mechanism. A next necessary step will be to add the finer and also critically important details of how NTD and CTD layers modulate the basic functions encoded in the receptor core.

## EXPERIMENTAL PROCEDURES

### Cells and Expression

Rat GluN1-1a (N1, U08261), GluN2A (N2A, M91561) or GluN2B (N2B, M91562) were expressed from pcDNA3.1(+) vectors in HEK293 cells along with GFP as described in detail previously (44).

NMDA receptor subunits lacking the NTD, N1^Δ^, N2A^Δ^, and N2B^Δ^, were constructed as follows. A plasmid encoding N1^Δ^ was previously developed by B. Laube (Darmstadt, Germany) and generously shared with us by P. Paoletti (Paris, France) (27). This construct contained the Kpnl-Sac1 fragment of the original GluN1-1a clone pN60 reported by the Nakanishi group (63), from which the nucleotide sequence encoding amino acids 5 to 358 was excised using PvuI. From this previously reported construct we sub-cloned the N1^Δ^ coding sequence into pcDNA 3.1 by engineering HindIII and Kozak sites just 5’ to the 22-residue signal peptide, and NotI sites just 3’ of the coding sequence. This manipulation added a consensus translation initiation site, preserved the signal peptide, and excluded a large 3’UTR from the initial construct, thus improving the efficiency of translation and surface expression in our HEK293 cell expression system.

The coding sequence for N2A^Δ^ in pCI was a gift from F. Zheng (Little Rock, AR). This cDNA originated from the EcoR I-XbaI fragment of pRSCI-A’ΔN1-3 constructed in the Neyton lab (24) which was reintroduced into wild-type N2A coding sequence with a XbaI site engineered to the pre-M1 region in the Zheng lab (64). The resulting construct preserved the 28-residue signal peptide, and excluded residues 5 - 385, such that the remaining coding sequence was identical to wild-type N2A. From this construct, we shuttled the EcoR1-Not1 fragment into pcDNA3.1.

We prepared the N2B^Δ^ coding plasmid by using the N2A^Δ^ (pcDNA3.1) described above, and replacing the fragment between the SacII site immediately upstream of the 27-residue signal peptide and immediately downstream of M386, with the corresponding SacII/NotI cassette of 2B^Δ^.

### Electrophysiology

*Whole-cell currents* were recorded with borosilicate glass electrodes filled with intracellular solution (in mM) 135 CsOH, 33 CsF, 2 MgCl_2_, 1 CaCl_2_, 10 HEPES and 11 EGTA, adjusted to pH 7.4 (CsOH). Cells were clamped at −70 mV and were perfused with extracellular solution containing (in mM) 150 NaCl, 2.5 KCl, 0.5 CaCl_2_, 10 HEPBS, 0.01 EDTA and 0.1 glycine, adjusted to pH 8 (NaOH). Responses were elicited with external solutions with added glutamate (1 mM). For zinc inhibition curves, extracellular solutions did not contain EDTA; free zinc concentrations in 10 mM tricine-buffered solutions were calculated using Maxchelator software (www.stanford.edu/ ~cpatton/ maxc.html) using a binding constant of 10^−5^ M as previously reported (37). Currents were amplified and low-pass filtered (2 kHz, Axopatch 200B), digitally sampled (5 kHz, Digidata 1440A) and acquired into digital files with pClamp 10.2 software. Traces were analyzed in Clampfit 10.2 and further with OriginPro8.

### Single-channel currents

were recorded continuously from cell-attached patches containing only one active receptor, with borosilicate electrodes filled with extracellular solution (in mM) 150 NaCl, 2.5 KCl, 10 HEPBS, 1 EDTA, 1 glutamate and 0.1 glycine, adjusted to pH 8 (NaOH). Inward Na^+^ currents were elicited by applying +100 mV through the recording pipette. Currents were amplified and low-pass filtered (10 kHz, Axopatch 200B), digitally sampled (40 kHz, National Instruments PCI-6229 A/D board) and acquired into digital files using QuB acquisition software (www.qub.buffalo.edu, University at Buffalo, Buffalo, NY).

### Kinetic Modeling

Selection, processing, idealization and modeling of microscopic data were done in QuB as described previously in detail (65). Briefly, processing and analyses were done on records selected to have only one active channel, which required minimum corrections and contained 3.0 x 10^4^ – 7.2 x 10^5^ events, after imposing a 0.075 ms resolution. Idealization (SKM algorithm) and modeling (MIL algorithm) were done in QuB using 12 kHz digitally filtered data (66,67). State models were fit to individual recordings by adding closed and open states, sequentially; best fitting models were selected with an arbitrarily set threshold of 10 log-likelihood units. Time constants and areas of individual kinetic components, as well as rate constants for the transitions considered were calculated for each data file with the models indicated and presented in Figures 3 and 4 as rounded means.

### Simulations

Macroscopic traces were simulated in QuB or MATLAB software, in response to square jumps into 1 mM glutamate, as the sum of time-dependent open state occupancies, using the experimentally determined microscopic rate constants for the best fitting models. Previously reported microscopic rates for glutamate binding and dissociation rate constants of wild-type receptors were used: 2 x 10^7^ M^−1^s^−1^ and 60 s^−1^, for N1/N2A (68) and 6 x 10^6^ M^−1^s^−1^ and 15 s^−1^, for N1/N2B (1,46,50). Simulated traces were analyzed in a manner similar to experimental macroscopic traces.

### Statistics

Results are reported for each data set as means ± SEM. Statistical differences were evaluated using two-tailed Student’s *t*-tests assuming equal variance, and were considered significant for p < 0.05.

## Acknowledgements

We thank Drs. Bodo Laube, Fang Zheng, Vasanthi Jayaraman, Pierre Paoletti and Jacques Neyton for sharing reagents; funding was from NIH DS052669 to GKP.

## FOOTNOTES

### Abbreviations used

NMDA: (N-methyl-D-aspartate) receptor
iGluR: ionotropic glutamate receptor
NTD: N-terminal domain
LBD: ligand binding domain
TMD: transmembrane domain
CTD: C-terminal domain
P_o_: open probability
MOT: mean open time
MCT: mean closed time
I_ss_: steady state current
I_pk_: peak current
τd: deactivation time constant
τ_D_: desensitization time constant

## REFERENCES

1. Iacobucci, G. J., and Popescu, G. K. (2017) NMDA receptors: linking physiological output to biophysical operation. Nat Rev Neurosci 18, 236–249

2. Popescu, G. K. (2012) Modes of glutamate receptor gating. J Physiol 590, 73–91

3. Hansen, K. B., Yi, F., Perszyk, R. E., Furukawa, H., Wollmuth, L. P., Gibb, A. J., and Traynelis, S. F. (2018) Structure, function, and allosteric modulation of NMDA receptors. J Gen Physiol 150, 1081–1105

4. Lu, W., Du, J., Goehring, A., and Gouaux, E. (2017) Cryo-EM structures of the triheteromeric NMDA receptor and its allosteric modulation. Science 355

5. Zhu, S., Stein, R. A., Yoshioka, C., Lee, C. H., Goehring, A., McHaourab, H. S., and Gouaux, E. (2016) Mechanism of NMDA Receptor Inhibition and Activation. Cell 165, 704–714

6. Tajima, N., Karakas, E., Grant, T., Simorowski, N., Diaz-Avalos, R., Grigorieff, N., and Furukawa, H. (2016) Activation of NMDA receptors and the mechanism of inhibition by ifenprodil. Nature 534, 63–68

7. Song, X., Jensen, M. Ø., Jogini, V., Stein, R. A., Lee, C.-H., McHaourab, H. S., Shaw, D. E., and Gouaux, E. (2018) Mechanism of NMDA receptor channel block by MK-801 and memantine. Nature 556, 515–519

8. Regan, M. C., Grant, T., McDaniel, M. J., Karakas, E., Zhang, J., Traynelis, S. F., Grigorieff, N., and Furukawa, H. (2018) Structural Mechanism of Functional Modulation by Gene Splicing in NMDA Receptors. Neuron 98, 521–529 e523

9. Chou, T.-H., Tajima, N., Romero-Hernandez, A., and Furukawa, H. (2020) Structural Basis of Functional Transitions in Mammalian NMDA Receptors. Cell

10. Traynelis, S. F., Wollmuth, L. P., McBain, C. J., Menniti, F. S., Vance, K. M., Ogden, K. K., Hansen, K. B., Yuan, H., Myers, S. J., and Dingledine, R. (2010) Glutamate receptor ion channels: structure, regulation, and function. Pharmacol Rev 62, 405–496

11. Nakanishi, N., Shneider, N. A., and Axel, R. (1990) A family of glutamate receptor genes: evidence for the formation of heteromultimeric receptors with distinct channel properties. Neuron 5, 569–581

12. O’Hara, P. J., Sheppard, P. O., Thogersen, H., Venezia, D., Haldeman, B. A., McGrane, V., Houamed, K. M., Thomsen, C., Gilbert, T. L., and Mulvihill, E. R. (1993) The ligand-binding domain in metabotropic glutamate receptors is related to bacterial periplasmic binding proteins. Neuron 11, 41–52

13. Stern-Bach, Y., Bettler, B., Hartley, M., Sheppard, P. O., O’Hara, P. J., and Heinemann, S. F. (1994) Agonist selectivity of glutamate receptors is specified by two domains structurally related to bacterial amino acid-binding proteins. Neuron 13, 1345–1357

14. Wo, Z. G., and Oswald, R. E. (1995) Unraveling the modular design of glutamate-gated ion channels. Trends Neurosci 18, 161–168

15. Wood, M. W., VanDongen, H. M., and VanDongen, A. M. (1995) Structural conservation of ion conduction pathways in K channels and glutamate receptors. Proc Natl Acad Sci U S A 92, 4882–4886

16. Armstrong, N., Sun, Y., Chen, G. Q., and Gouaux, E. (1998) Structure of a glutamate-receptor ligand-binding core in complex with kainate. Nature 395, 913–917

17. Hirai, H., Kirsch, J., Laube, B., Betz, H., and Kuhse, J. (1996) The glycine binding site of the N-methyl-D-aspartate receptor subunit NR1: identification of novel determinants of co-agonist potentiation in the extracellular M3-M4 loop region. Proc Natl Acad Sci U S A 93, 6031–6036

18. Laube, B., Hirai, H., Sturgess, M., Betz, H., and Kuhse, J. (1997) Molecular determinants of agonist discrimination by NMDA receptor subunits: analysis of the glutamate binding site on the NR2B subunit. Neuron 18, 493–503

19. Verdoorn, T. A., Burnashev, N., Monyer, H., Seeburg, P. H., and Sakmann, B. (1991) Structural determinants of ion flow through recombinant glutamate receptor channels. Science 252, 1715–1718

20. Burnashev, N., Schoepfer, R., Monyer, H., Ruppersberg, J. P., Gunther, W., Seeburg, P. H., and Sakmann, B. (1992) Control by asparagine residues of calcium permeability and magnesium blockade in the NMDA receptor. Science 257, 1415–1419

21. McDaniel, M. J., Ogden, K. K., Kell, S. A., Burger, P. B., Liotta, D. C., and Traynelis, S. F. (2020) NMDA receptor channel gating control by the pre-M1 helix. J Gen Physiol 152

22. Iacobucci, G. J., Wen, H., Helou, M. B., Zheng, W., and Popescu, G. K. (2020) Cross-subunit Interactions that Stabilize Open States Mediate Gating in NMDA Receptors. bioRxiv, 2020.2006.2008.140525

23. Ladislav, M., Cerny, J., Krusek, J., Horak, M., Balik, A., and Vyklicky, L. (2018) The LILI Motif of M3-S2 Linkers Is a Component of the NMDA Receptor Channel Gate. Front Mol Neurosci 11, 113

24. Fayyazuddin, A., Villarroel, A., Le Goff, A., Lerma, J., and Neyton, J. (2000) Four residues of the extracellular N-terminal domain of the NR2A subunit control high-affinity Zn2+ binding to NMDA receptors. Neuron 25, 683–694

25. Meddows, E., Le Bourdelles, B., Grimwood, S., Wafford, K., Sandhu, S., Whiting, P., and McIlhinney, R. A. (2001) Identification of molecular determinants that are important in the assembly of NMDA receptors. J Biol Chem 276, 18795–18803

26. Rachline, J., Perin-Dureau, F., Le Goff, A., Neyton, J., and Paoletti, P. (2005) The Micromolar Zinc-Binding Domain on the NMDA Receptor Subunit NR2B. J. Neurosci. 25, 308–317

27. Madry, C., Mesic, I., Betz, H., and Laube, B. (2007) The N-terminal domains of both NR1 and NR2 subunits determine allosteric Zn2+ inhibition and glycine affinity of NMDA receptors. Mol Pharmacol, mol.107.040071

28. Gielen, M., Siegler Retchless, B., Mony, L., Johnson, J. W., and Paoletti, P. (2009) Mechanism of differential control of NMDA receptor activity by NR2 subunits. Nature 459, 703–707

29. Yuan, H., Hansen, K. B., Vance, K. M., Ogden, K. K., and Traynelis, S. F. (2009) Control of NMDA receptor function by the NR2 subunit amino-terminal domain. J Neurosci 29, 12045–12058

30. Yuan, H., Vance, K. M., Junge, C. E., Geballe, M. T., Snyder, J. P., Hepler, J. R., Yepes, M., Low, C. M., and Traynelis, S. F. (2009) The serine protease plasmin cleaves the amino-terminal domain of the NR2A subunit to relieve zinc inhibition of the NMDA receptors. J Biol Chem 284, 12862–12873

31. Maki, B. A., Aman, T. K., Amico-Ruvio, S. A., Kussius, C. L., and Popescu, G. K. (2012) C-terminal domains of NMDA acid receptor modulate unitary channel conductance and gating. J Biol Chem 287, 36071–36080

32. Chen, G. Q., Cui, C., Mayer, M. L., and Gouaux, E. (1999) Functional characterization of a potassium-selective prokaryotic glutamate receptor. Nature 402, 817–821

33. Iacobucci, G. J., and Popescu, G. K. (2019) N-Methyl-D-Aspartate Receptors. in Oxford Handbooks Online (Bhattacharjee, A. ed.), Oxford University Press. pp

34. Furukawa, H. (2011) Structure and function of glutamate receptor amino terminal domains. The Journal of Physiology

35. Paoletti, P., Bellone, C., and Zhou, Q. (2013) NMDA receptor subunit diversity: impact on receptor properties, synaptic plasticity and disease. Nat Rev Neurosci 14, 383–400

36. Paoletti, P. (2011) Molecular basis of NMDA receptor functional diversity. Eur J Neurosci 33, 1351–1365

37. Paoletti, P., Ascher, P., and Neyton, J. (1997) High-affinity zinc inhibition of NMDA NR1-NR2A receptors. J Neurosci 17, 5711–5725

38. Paoletti, P., Perin-Dureau, F., Fayyazuddin, A., Le Goff, A., Callebaut, I., and Neyton, J. (2000) Molecular organization of a zinc binding N-terminal modulatory domain in a NMDA receptor subunit. Neuron 28, 911–925

39. Christine, C. W., and Choi, D. W. (1990) Effect of zinc on NMDA receptor-mediated channel currents in cortical neurons. J Neurosci 10, 108–116

40. Williams, K. (1996) Separating dual effects of zinc at recombinant N-methyl-D-aspartate receptors. Neurosci Lett 215, 9–12

41. Chen, N., Moshaver, A., and Raymond, L. A. (1997) Differential sensitivity of recombinant NMDA receptor subtypes to zinc inhibition. Mol Pharmacol 51, 1015–1023

42. Howe, J. R., Colquhoun, D., and Cull-Candy, S. G. (1988) On the kinetics of large-conductance glutamate-receptor ion channels in rat cerebellar granule neurons. Proc R Soc Lond B Biol Sci 233, 407–422

43. Nowak, L., Bregestovski, P., Ascher, P., Herbet, A., and Prochiantz, A. (1984) Magnesium gates glutamate-activated channels in mouse central neurones Nature 307, 462–465

44. Kussius, C. L., Kaur, N., and Popescu, G. K. (2009) Pregnanolone sulfate promotes desensitization of activated NMDA receptors. J Neurosci 29, 6819–6827

45. Borschel, W. F., Myers, J. M., Kasperek, E. M., Smith, T. P., Graziane, N. M., Nowak, L. M., and Popescu, G. K. (2012) Gating reaction mechanism of neuronal NMDA receptors. J Neurophysiol 108, 3105–3115

46. Amico-Ruvio, S. A., and Popescu, G. K. (2010) Stationary gating of GluN1/GluN2B receptors in intact membrane patches. Biophys J 98, 1160–1169

47. Banke, T. G., and Traynelis, S. F. (2003) Activation of NR1/NR2B NMDA receptors. Nat Neurosci 6, 144–152

48. Popescu, G., and Auerbach, A. (2003) Modal gating of NMDA receptors and the shape of their synaptic response. Nat Neurosci 6, 476–483

49. Auerbach, A., and Zhou, Y. (2005) Gating reaction mechanisms for NMDA receptor channels. J Neurosci 25, 7914–7923

50. Iacobucci, G. J., and Popescu, G. K. (2018) Kinetic models for activation and modulation of NMDA receptor subtypes. Current Opinion in Physiology 2, 114–122

51. Carmignoto, G., and Vicini, S. (1992) Activity-dependent decrease in NMDA receptor responses during development of the visual cortex. Science 258, 1007–1011

52. Cline, H. T., Debski, E. A., and Constantine-Paton, M. (1990) The role of the NMDA receptor in the development of the frog visual system. Adv Exp Med Biol 268, 197–203

53. Williams, K., Russell, S. L., Shen, Y. M., and Molinoff, P. B. (1993) Developmental switch in the expression of NMDA receptors occurs in vivo and in vitro. Neuron 10, 267–278

54. Groc, L., Heine, M., Cousins, S. L., Stephenson, F. A., Lounis, B., Cognet, L., and Choquet, D. (2006) NMDA receptor surface mobility depends on NR2A-2B subunits. Proc Natl Acad Sci U S A 103, 18769–18774

55. Vicini, S., Wang, J. F., Li, J. H., Zhu, W. J., Wang, Y. H., Luo, J. H., Wolfe, B. B., and Grayson, D. R. (1998) Functional and pharmacological differences between recombinant N-methyl-D-aspartate receptors. in J Neurophysiol

56. Monyer, H., Sprengel, R., Schoepfer, R., Herb, A., Higuchi, M., Lomeli, H., Burnashev, N., Sakmann, B., and Seeburg, P. H. (1992) Heteromeric NMDA receptors: molecular and functional distinction of subtypes. Science 256, 1217–1221

57. Furukawa, H., and Gouaux, E. (2003) Mechanisms of activation, inhibition and specificity: crystal structures of the NMDA receptor NR1 ligand-binding core. Embo J 22, 2873–2885

58. Karakas, E., and Furukawa, H. (2014) Crystal structure of a heterotetrameric NMDA receptor ion channel. Science 344, 992–997

59. Chou, T. H., Tajima, N., Romero-Hernandez, A., and Furukawa, H. (2020) Structural Basis of Functional Transitions in Mammalian NMDA Receptors. Cell 182, 357–371 e313

60. Maki, B. A., Cole, R., and Popescu, G. K. (2013) Two serine residues on GluN2A C-terminal tails control NMDA receptor current decay times. Channels (Austin) 7, 126–132

61. Aman, T. K., Maki, B. A., Ruffino, T. J., Kasperek, E. M., and Popescu, G. K. (2014) Separate intramolecular targets for protein kinase A control NMDA receptor gating and Ca2+ permeability. J Biol Chem 289, 18805–18817

62. Lee, C. H., Lu, W., Michel, J. C., Goehring, A., Du, J., Song, X., and Gouaux, E. (2014) NMDA receptor structures reveal subunit arrangement and pore architecture. Nature 511, 191–197

63. Moriyoshi, K., Masu, M., Ishii, T., Shigemoto, R., Mizuno, N., and Nakanishi, S. (1991) Molecular cloning and characterization of the rat NMDA receptor. Nature 354, 31–37

64. Hu, B., and Zheng, F. (2005) Molecular Determinants of Glycine-independent Desensitization of NR1/NR2A Receptors. J Pharmacol Exp Ther, jpet.104.080168

65. Cummings, K., Iacobucci, G., and Popescu, G. (2016) Extracting Rate Constants for NMDA Receptor Gating from One-Channel Current Recordings. in Ionotropic Glutamate Receptor Technologies (Popescu, G. K. ed.), Springer New York. pp 273–299

66. Qin, F., Auerbach, A., and Sachs, F. (1997) Maximum likelihood estimation of aggregated Markov processes. Proc Biol Sci 264, 375–383

67. Qin, F., Auerbach, A., and Sachs, F. (2000) Hidden Markov modeling for single channel kinetics with filtering and correlated noise. Biophys J 79, 1928–1944.

68. Popescu, G., Robert, A., Howe, J. R., and Auerbach, A. (2004) Reaction mechanism determines NMDA receptor response to repetitive stimulation. Nature 430, 790–793

